# Enhancing altruism by electrically augmenting frontoparietal gamma-band phase coupling

**DOI:** 10.1101/2024.04.01.587528

**Authors:** Jie Hu, Marius Moisa, Christian C. Ruff

## Abstract

Cooperation, productivity, and cohesion in human societies depend on altruism, the tendency to share resources with others even though this is costly. While altruism is a widely shared social norm, people vary strongly in their inclination to behave altruistically, in particular across situations with different types of inequality in resource distribution. What neurobiological factors underlie this variability? And can these be targeted by interventions to enhance altruistic behavior? Here, we build on recent EEG findings that altruistic choices during disadvantageous inequality correlate with oscillatory gamma-band coherence between frontal regions (representing other’s interest) and parietal regions (representing neural evidence accumulation). We apply a transcranial alternating current stimulation (tACS) protocol designed to exogenously enhance this fronto-parietal coherence and find that this leads to increased altruism, specifically during disadvantageous inequality as hypothesized based on the EGG findings. Computational modeling reveals that this transcranial entrainment does not just add noise to the decision process but specifically increases the weight individuals assign to other-regarding concerns during choices. Our findings show that altruism can be enhanced by neurostimulation designed to enhance oscillatory synchronization between frontal and parietal areas. This establishes a neural basis for altruism and identifies a neural target for interventions aimed at improving prosocial behavior.

## Introduction

Altruism is the foundation for collaboration and solidarity in human society (de Waal, 2008; Simpson & Willer, 2015). Lacking altruism is a hallmark of social deficits in psychiatric and neurological disorders (e.g. psychopathy, autism, and alexithymia)(Bird & Viding, 2014; FeldmanHall et al., 2013) and lies at the heart of many social problems at the community level (e.g. high crime rate and poor crisis management)(Chamlin & Cochran, 1997; Rose et al., 2022). The neural basis of altruism is therefore intensely studied in neuroscience, with the aim to identify neural mechanisms that may be targeted by interventions to enhance altruistic behavior.

Previous studies on this topic have shown that altruistic decisions involve separable neural representations of the choice outcomes for oneself versus the other person, as well as processes that integrate these two types of choice evidence (Crockett et al., 2017; Gao et al., 2018; Hutcherson et al., 2015; Konovalov et al., 2021). Most of these studies have focused on characterizing the precise roles of individual brain regions – for examples, some studies have shown that activity in ventro- and dorsomedial prefrontal cortex (vmPFC/dmPFC) represents the size of potential monetary outcomes for others vs oneself (FeldmanHall et al., 2015; Hu et al., 2021; Lockwood et al., 2015), whereas parietal areas (e.g., temporoparietal junction - TPJ and inferior parietal cortex) appear to encode signals that compare these magnitudes (e.g., equality) or that represent categorizations of the competing choice options (e.g. as selfish or altruistic) (Hutcherson et al., 2015; Morishima et al., 2012). However, how all these different signals may be integrated at the neural level to determine the choice outcome has only been examined by a few studies (Hare et al., 2010; Hu et al., 2017). As an example, one study has shown that communication between brain areas may play a role, since altruistic donations were accompanied by stronger connectivity between frontal (e.g. vmPFC) and parietal cortices (e.g. TPJ) (Hare et al., 2010).

What specific functional role may frontoparietal interactions play for altruistic decision-making? One possible mechanism is information exchange by oscillatory synchronization, as suggested by findings in the non-social decision-making domain: During choices between food items, the evidence for a particular choice option appears to be determined by neural value computations and comparisons in medial prefrontal cortex, which are then transformed into response signals in the parietal cortex via oscillatory gamma-band phase-coupling between these two regions (Basten et al., 2010; Pisauro et al., 2017; Polanía et al., 2014, 2015). Similar mechanisms may also be important for altruistic choice: A recent electroencephalography (EEG) study showed that the strength of fronto-parietal gamma-band phase-coupling is positively associated with individuals’ altruistic preferences (Hu et al., 2023). Since interregional gamma-band synchronisation is thought to facilitate that coordinated activity in one neural population impacts maximally on the activity in interconnected neural populations (Siegel et al., 2012; Thut et al., 2012), the fronto-parietal phase-coupling observed in the study may serve to allow other-regarding concerns (as represented in frontal cortex) to impact on parietal evidence integration processes guiding choice (Brosnan et al., 2020; Hu et al., 2023; Polanía et al., 2014). However, previous neuroimaging studies could only provide correlative evidence for this possible relationship between frontoparietal interaction and altruism (Edelson et al., 2018; Hare et al., 2010a; Harris et al., 2018; Hu et al., 2017). Experimentally clarifying the causal role of frontoparietal connectivity for altruism may be a necessary first step towards developing and/or improving intervention approaches to facilitate altruism, e.g., in people affected by social apathy and other related psychiatric and neurological disorders.

One factor that complicates the study of the neural basis of altruism is that altruistic decisions can be driven by distinct motives in different contexts(Hein et al., 2016; Hu et al., 2023; Piliavin & Charng, 1990; Zaki & Mitchell, 2011). For instance, any inequality in resources between two individuals can substantially affect wealth distribution choices: When people possess more than others (advantageous inequality, ADV), stronger “guilt” emotion may drive them to share more; whereas when people possess less than others (disadvantageous inequality, DIS), “envy” emotion may drive them to share less (Charness & Rabin, 2002; Fehr & Schmidt, 1999; Gao et al., 2018; Güroğlu et al., 2014; Morishima et al., 2012; Tricomi et al., 2010). While in both these contexts, similar parietal evidence accumulation signals could be observed in the EEG signal, the strength of frontoparietal phase-coupling was associated with altruistic preferences mainly during disadvantageous inequality (Hu et al., 2023). Irrespective of these contextual differences, it is generally unclear whether the frontoparietal oscillatory coherence evident in the EEG signal is linked to altruism in a causal sense - either by implementing a general tendency or a specific motive underlying altruism in a specific context.

In the current study, we address these issues by combining a modified dictator game with a high-definition (HD) transcranial alternating current stimulation (tACS) approach designed to exogenously modulate phase coupling of distant cortical brain regions (Saturnino et al., 2017; Violante et al., 2017). This approach allows us to clarify three fundamental questions regarding the role of interregional phase-coupling in altruistic decision-making. First: Is it in principle possible to enhance altruism by external stimulation designed to affect neural coherence? Second: Is it specifically gamma-band frontoparietal synchronization that is causally relevant for altruistic choices? Third: Is gamma-band frontoparietal synchronization relevant for altruistic motivation in general, or just for specific motives present during disadvantageous (versus advantageous) inequality?

## Results

Participants played as proposers in a modified Dictator Game and had to choose between two possible monetary distributions between themselves and anonymous partners (Figure 1A). As in our previous study, we varied the sizes of payoffs in a way that they created two different inequality contexts – disadvantageous (DIS) in which participants got less than their partners for both distribution options and advantageous (ADV) in which participants got more than their partners for both distribution options (for detailed trial set description see Figure S1). These two types of trials were only defined by the size of the payoffs presented on the screen (i.e., there were no other visual markers) and were randomly intermixed in a manner that led to strongly varying inequality and context variation from trial to trial, while still ensuring an overall balanced presentation across the whole experiment (see Figure S1).

During the task, participants received transcranial alternative current stimulation (tACS) by means of a high-definition tACS setup designed to non-invasively synchronise cortical rhythms at a specific frequency between the frontal and parietal regions underlying the electrodes (Saturnino et al., 2017; Violante et al., 2017). To this end, we placed two arrays of small electrodes (3 × 1 HD electrode montage) over the specific positions where we had previously observed the coherence pattern (Hu et al., 2023) and provided focal electrical current stimulation to entrain the coherence between the corresponding frontal and parietal areas (Figure 1B). Participants received three different types of entrainment: gamma, alpha, and sham stimulation (Figure 1C&D). Importantly, the tACS oscillations were temporally aligned between the two arrays of electrodes (i.e., in-phase stimulation, phase = 0). We included alpha entrainment as an active control condition for potentially non-specific effects of the stimulation (Soutschek et al., 2021). Since our previous study had found that greater altruistic preferences correlate with stronger frontoparietal synchronization in the gamma-band frequency (Hu et al., 2023), we expected that only gamma, but not alpha entrainment, would enhance altruistic preferences. For the sham stimulation, we provided either gamma (for the sham block following the gamma-entrainment block) or alpha entrainment (for the sham block following the alpha-entrainment block) for only 2 s every 27 s during the two sham stimulation blocks within each experimental run. The order of the entrainment type (i.e., gamma or alpha) for the two sham stimulation blocks was counterbalanced across different experimental runs (see STAR Methods section for more details).

**Figure 1.**
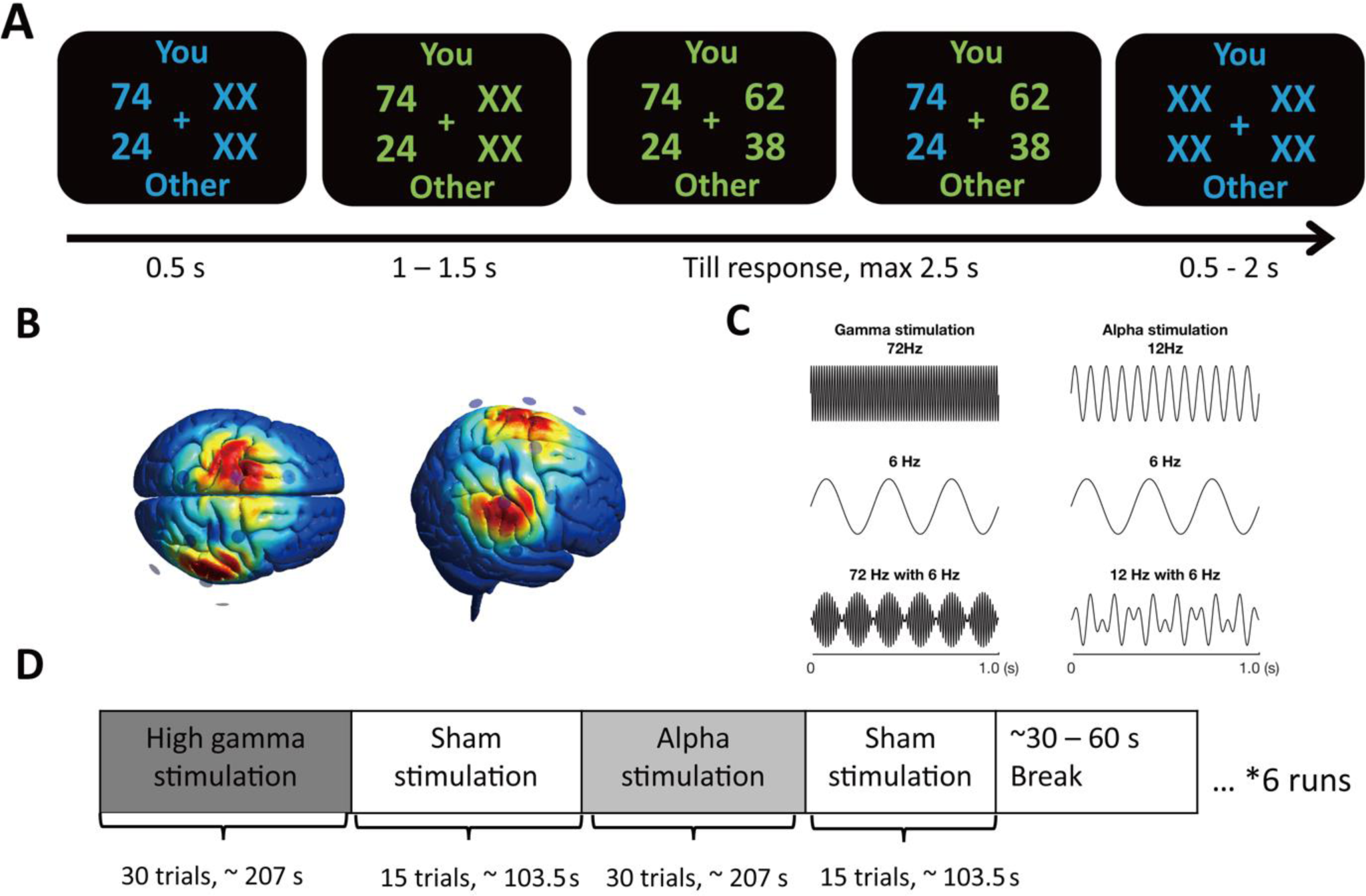
Experimental design and protocol. (A) Example of a single trial. Participants acted as proposers to distribute a certain amount of monetary tokens between themselves and anonymous partners. In the beginning of each trial, participants were presented with one reference option in blue for 0.5 s. The trial starts when the font color changes from blue to green. After the presentation of the 2^nd^ option, participants needed to choose between the two options within 2.5 s. The selected option was highlighted in blue before the inter-trial interval. Font color assignment to phases (i.e., blue and green to response) was counterbalanced across participants. (B) tACS electrode montage and electric field density simulation. Two sets of small electrodes (3 ×1 HD electrode montage) were mounted over frontal and parietal regions which were located based on our previous EEG study (Hu et al., 2022).(Hu et al., 2022).(Hu et al., 2022). The normalized simulated electric field distribution showed that the targeted parietal and frontal areas under the two sets of tACS electrodes are affected by the stimulation with a relatively good special focality. (C) Schematic of the alternating currents delivered to participants. Participants received alternating currents at 72 Hz with an envelop of 6 Hz (gamma-band stimulation, left bottom panel), alternating currents at 12 Hz with an envelop of 6 Hz (alpha-band stimulation, right bottom panel) as active control stimulation, and sham stimulation. (D) Participants received these three types of stimulation during the task. Each run lasted approx. 7 min and the order of the stimulation blocks was counterbalanced across different runs.

To examine how individuals’ altruistic behavior was influenced by the different types of stimulation, we performed linear mixed-effects regression of altruistic choice (more precisely, the probability of choosing the more altruistic option on each trial) on inequality context (ADV vs. DIS), stimulation type (gamma vs. alpha or gamma vs. sham), and their interactions. In line with our hypothesis, we found that gamma entrainment indeed led to an increase in the probability of choosing the altruistic option, compared to both alpha entrainment (main effect of gamma entrainment in Table 1 Model 1: *β* = 0.18 ± 0.10, Estimate ± SE, *p* = 0.032) and sham stimulation (main effect of gamma entrainment in Table 1 Model 2: *β* = 0.19 ±0.10, *p* = 0.028). Altruistic choice probability did not differ between alpha entrainment and sham stimulation [main effects of sham (alpha) stimulation in Table 1 Model 1 (Model 2): *β* = −0.01 ±0.10 (*β* = −0.01 ±0.10), *p* = 0.942]. This shows that fronto-parietal gamma-band coherence is indeed causally relevant for altruistic behavior, and that it is possible to enhance altruism by strengthening this coherence.

As in our previous EEG study, the probability of altruistic choice was higher in the ADV than in the DIS context (main effect of inequality context in Table 1 Model 1: *β* = 0.82 ±0.09, *p* < 0.001; in Table 1 Model 2: *β* = 0.89 ±0.09, *p* < 0.001), consistent with previous suggestions that individuals consider others’ interest more in their choices when they possess more compared to when they possess less than others (Hu et al., 2023). As we stimulated each participant with an electric current intensity in line with his/her tolerance level for the stimulation currents tested before the experiment runs, the peak-to-peak current intensity for different participants varied between 2.4 mA and 4 mA. Therefore, in the regression analyses, we also included individual electric current intensity for each participant and individual discomfort ratings for each stimulation run as control variables. The effects reported above were not influenced by potential differences in discomfort experienced by the participants for different types of entrainment, or by the individual electric current intensity the participants received (Table 1 Model 3 & 4, see also Methods).

Although the interaction between stimulation type (gamma vs alpha or gamma vs. sham) and inequality context was not statistically significant, inspection of the data in the two inequality contexts showed that the main effect of gamma entrainment appeared to mainly reflect the increasing effect of gamma entrainment on altruistic choice in the DIS context (gamma: 0.12 ± 0.02; versus sham: 0.10 ± 0.02, *t(43)* = 2.07, *p* = 0.044, *Cohen’s d* = 0.31; versus alpha: 0.10 ±0.02, *t(43)* = 2.27, *p* = 0.028, *Cohen’s d* = 0.34). In the ADV context, no such effects were evident (gamma: 0.20 ±0.03, alpha: 0.19 ±0.03, sham: 0.20 ±0.03, all *p*s > 0.09) (Figure 2A & B, Table S1 & S2). This suggests that fronto-parietal gamma-band coherence may not be equally important for all motives underlying altruistic choice, but may particularly affect those motives guiding choices during disadvantageous inequality.

**Table 1.**
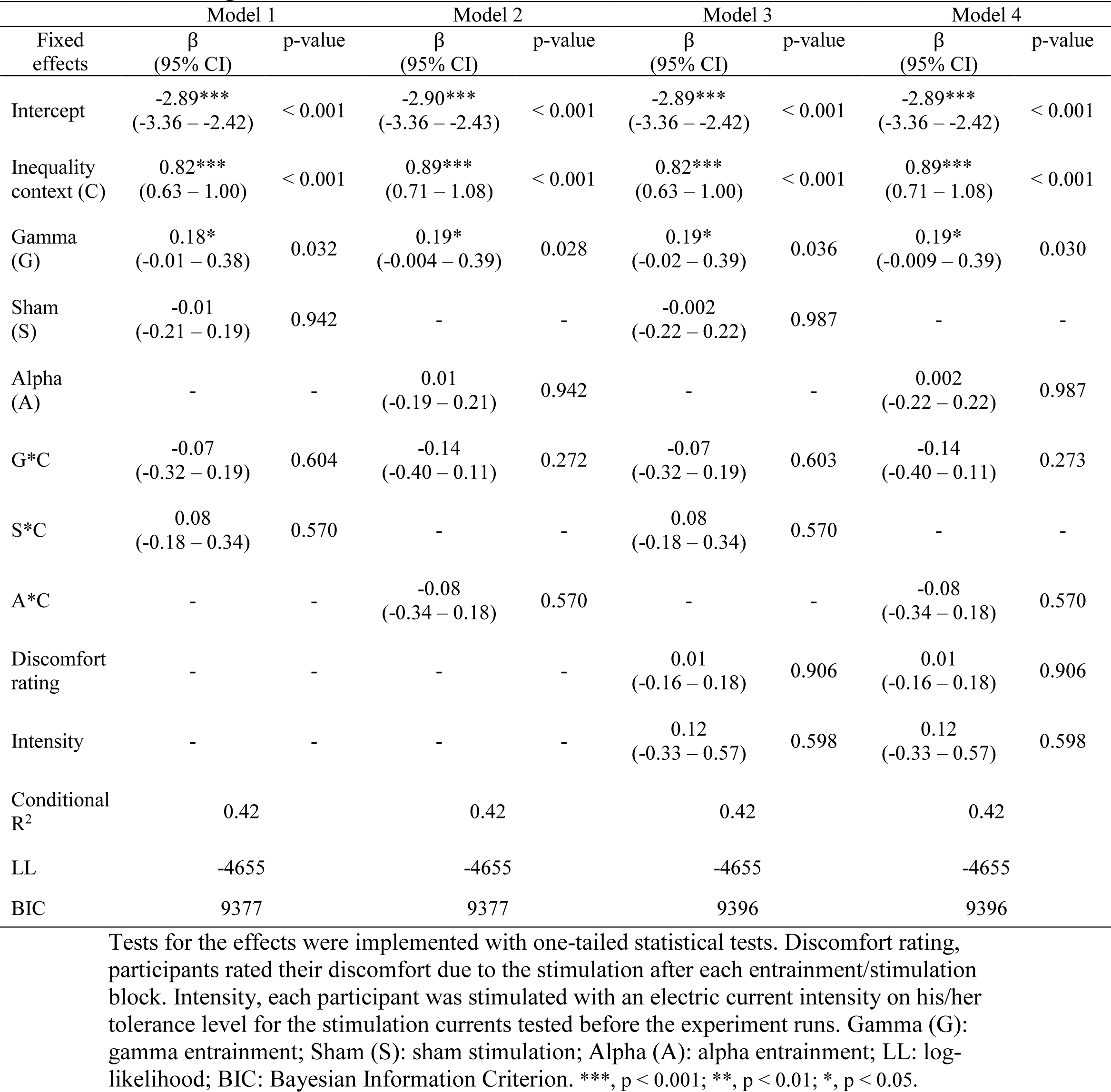
Logistic mixed-effects model results of choice data.

**Figure 2.**
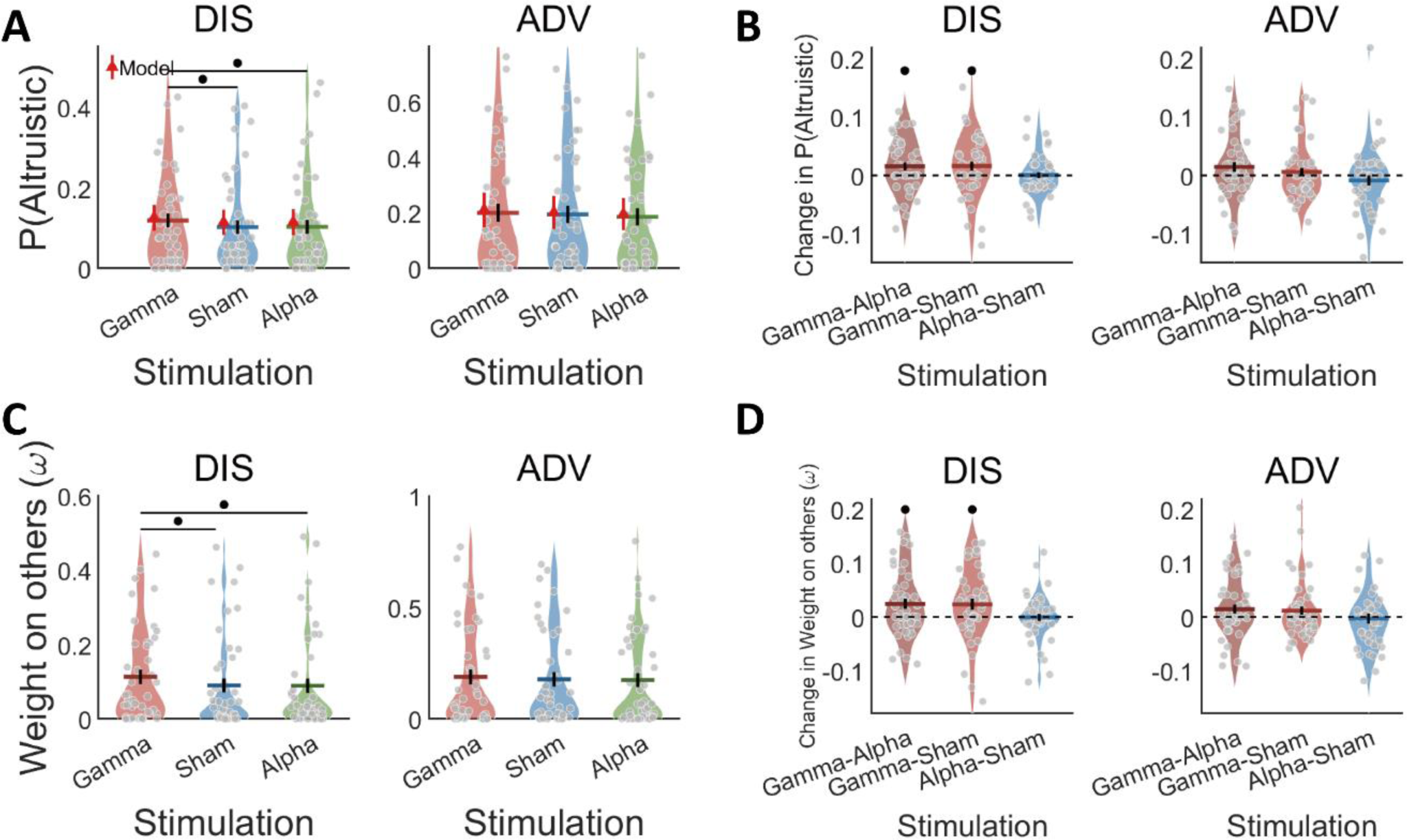
Behavioral and modelling results. (A) The probability of altruistic choice in DIS context was higher during gamma entrainment relative to both sham and alpha entrainment. The winning model recovers the effects of stimulation. Colored bars indicate the means and black error bars display the standard errors of the mean (SE). The red triangle dots and error bars represent the model simulation mean and 95% confidence intervals (CIs). (B) The difference in the probability of altruistic choice in the DIS context was significantly higher than 0 for gamma entrainment compared to both alpha entrainment and sham stimulation. (C) The winning model shows that gamma entrainment, relative to both sham and alpha entrainment, increased the weight on others in the DIS context. (D) The difference in the parameter of weight on others in the DIS context was significantly higher than 0 for gamma entrainment compared to both alpha entrainment and sham stimulation. Colored bars indicate the means for each stimulation type, and black error bars display the standard errors of the mean (SE). Each grey dot represents one participant. •, *p* < 0.05.

How exactly does gamma entrainment promote altruistic choice? In principle, altruistic choices could increase because the stimulation enhances the influence of other-regarding concerns on behavior, in line with the notion that the stimulation increases how behavior is guided by altruistic motives. Alternatively, the stimulation could impact on more general aspects of the choice process that nevertheless can alter the apparent relative impacts of selfish and other-regarding concerns (Hämmerer et al., 2016; Li et al., 2020). For instance, the stimulation could change the consistency of choices with (altruistic) decision values (Hämmerer et al., 2016), or it could alter the decision weights people place on the efficiency of the monetary distributions (i.e., on the overall benefit for both parties, irrespective of how this is distributed) (Li et al., 2020).

To test which of these possible effects was most evident in our data, we implemented computational modeling analyses. These examined with comparison of three alternative models how our tACS entrainment affected different latent model variables corresponding to these different aspects of the choice process. In all the alternative models, we captured how each individual weighed others’ interest versus their own interest to construct an integrated decision utility. To also assess consistency with the computed integrated decision values, we included either a single inverse temperature parameter in model M1 or condition-specific inverse temperature parameters in model M2. Finally, in a third model (M3) we additionally tested the possibility that individuals’ decisions may depend not only on their own and other’s interest but also on the overall benefit for both parties (i.e., efficiency).

Specifically, in M1, the subjective value difference between the more generous option and the more selfish option (VD) was constructed based on the Charness-Rabin utility model (Charness & Rabin, 2002) in which people weigh between outcomes for themselves and for others (for detailed model specification and construction, see STAR Methods section). We calculated the trial-by-trial value difference (VD) between the two options and employed a softmax function with a single inverse temperature parameter to transform these value differences into probabilities of choosing the more altruistic option in different conditions. In M2, we included six inverse temperature parameters for the six different conditions. In M3, we included a set of parameters to measure to what extent individuals’ decisions were also based on efficiency considerations.

Model comparison analysis showed that the first model (M1) outperformed the other models (Table S3). M1 had the lowest BIC value among the three models and the Bayes factors (BFs) relative to the alternative models were over 100, indicating “very strong” evidence that M1 is superior to the other models. Posterior predictive checks showed that the altruistic choice probabilities predicted by the fitted winning model indeed captured the observed choice probabilities well (Figure S2); moreover, the same logistic mixed-effects regression of these simulated choices revealed similar effects as participants’ real choices. Specifically, the probability of altruistic choice was increased by gamma entrainment, both relative to alpha entrainment (main effect of gamma entrainment in Table S4 Model 1: *β* = 0.23 ± 0.10, *p* = 0.008) and relative to sham stimulation (main effect of gamma entrainment in Table S4 Model 2: *β* = 0.24 ±0.10, *p* = 0.007). We also performed parameter recovery analyses to validate the winning model. The results show that we could reliably recover the parameters in M1 (Figure S3), confirming that the winning model captures the specific motives and choice processes underlying participants’ altruistic behavior.

To confirm that gamma-band frontoparietal phase-coupling indeed increased individuals’ altruistic preferences (i.e., their decision weight over others’ interest, ω), we analyzed how the tACS changed the winning model’s parameters. We did so with analysis of variance (ANOVA) in the M1 altruistic preference parameter ω, using tACS entrainment and inequality context as the independent variables. This confirmed that the stimulation changed altruistic preference, with a significant main effect of entrainment (*F*(2, 86) = 4.56, *p* = 0.013, 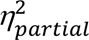 = 0.10) and a significant main effect of inequality context (*F*(1, 43) = 10.31, *p* = 0.003, 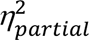 = 0.19), but an insignificant interaction effect between entrainment and inequality context (*F*(2, 86) = 0.72, *p* = 0.490, 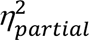 = 0.02). Post-hoc analyses confirmed that individuals’ altruistic preferences were increased by gamma entrainment (0.15 ± 0.02) relative to both alpha entrainment (0.13 ±0.02, *p* = 0.005, *Cohen’s d* = 0.41) and sham stimulation (0.13 ±0.02, *p* = 0.013, *Cohen’s d* = 0.37). Again, this effect was specific to gamma stimulation, as there were no significant differences in altruistic preferences between alpha entrainment (0.13 ±0.02) and sham stimulation (0.13 ±0.02, *p* = 0.386, *Cohen’s d* = 0.04). Finally, and again consistent with the model-free analyses, individuals’ altruistic preferences were stronger during advantageous (ADV, 0.18 ± 0.03) than disadvantageous inequality (DIS, 0.10 ±0.02, *p* = 0.002, *Cohen’s d* = 0.48), and only in the DIS context were the individuals’ weights on other interest significantly increased by gamma entrainment (0.11 ±0.02; both relative to sham (0.09 ±0.02), *t(43)* = 2.28, *p* = 0.042, *Cohen’s d* = 0.34 and to alpha (0.09 ± 0.02), *t(43)* = 2.59, *p* = 0.020, *Cohen’s d* = 0.39). The corresponding effects in the ADV context were small and all failed to reach significance (gamma entrainment: 0.19 ±0.03, alpha entrainment: 0.17 ±0.03, sham simulation: 0.18 ±0.03, *p*s > 0.1) (Figure 2C & D). The modeling analyses therefore confirmed that gamma entrainment indeed increased altruistic concerns, in a manner that was more marked during situations with disadvantageous inequality where participants had less than the other participant to begin with.

## Discussion

It is widely acknowledged that a lack of altruism leads to greater problems for individuals’ interpersonal interactions and for human society in general (Bird & Viding, 2014; Chamlin & Cochran, 1997; FeldmanHall et al., 2013; Rose et al., 2022). Therefore, clarifying the neural basis of altruism and developing efficient interventions to enhance altruism has become an important issue. Although previous studies have intensively explored putative neural mechanisms by which our brains may take altruistic decisions (e.g., specific patterns of brain activity and/or connectivity correlating with valuation and evidence accumulation (Hare et al., 2010; Hu et al., 2023; Hutcherson et al., 2015), there is still little evidence that such mechanisms are in fact causally relevant for altruistic behavior. This is particularly true for brain network mechanisms that subserve functional interactions between brain areas (Grover et al., 2021; Reinhart & Nguyen, 2019). To fill in this gap, our study provides evidence that altruism can be substantially enhanced by exogenously increasing gamma-band frontoparietal oscillatory synchronization.

Our results show that enhancing frontoparietal coherence in the gamma-band leads to more altruistic choices by strengthening altruistic preferences, suggesting that more efficient information sharing and integration of other-interest between the frontal and parietal cortex may promote more prosocial behavior. This interpretation is consistent with findings that frontal EEG responses to other-interest are correlated with individuals’ altruistic preferences (Hu et al., 2023), and with the well-documented role of parietal regions in evidence accumulation processes underlying nonsocial (Brosnan et al., 2020; Kelly & O’Connell, 2013) and social decision making (Hu et al., 2023; Pisauro et al., 2017; Polanía et al., 2014; Yao et al., 2020; Zhong et al., 2019; Zhou & Freedman, 2019). Our causal results integrate these separate sets of observations and suggest that frontal regions implement signals that evaluate others’ interests, which are communicated to parietal regions where evidence is integrated to guide the appropriate decisions. Importantly, such a mechanistic process is consistent with empirical patterns of both functional (Basten et al., 2010; Pisauro et al., 2017) and anatomical connectivity (Brosnan et al., 2020) between frontal and parietal regions. Our results therefore provide causal evidence for proposals that frontoparietal circuitry serves as a neural pathway for information sharing and integration underlying decision-making (Hare et al., 2011; Pisauro et al., 2017; Polanía et al., 2014, 2015), and establish that this general pathway also underlies altruistic choice.

As a difference to previous findings, we found that tACS designed to increase frontoparietal oscillatory coherence changed the decision weight over specific choice attributes (others payoffs), rather than choice variability as observed previously for non-social decision-making (Polanía et al., 2015). A potential explanation for this discrepancy is that the frontal signals targeted in the two studies may have originated in different frontal regions, which may play different roles for non-social versus altruistic decision-making. The signals/electrode positions identified and stimulated during non-social decision-making (Polanía et al., 2015) corresponded to the frontal pole, whereas the signals/electrode positions targeted here to increase altruism corresponded to a relatively more posterior frontal region (Hu et al., 2023). In other brain stimulation studies of social decision making, yet other specific prefrontal areas were targeted based on different anatomical locations identified with neuroimaging (Knoch et al., 2006, 2008, 2009; Ruff et al., 2013). While the functional resolution of brain stimulation methods is obviously limited (Polanía et al., 2018), our results clearly warrant that future studies should try to differentiate different fronto-parietal circuits and characterize how oscillatory coherence within them affects different types of decision-making (e.g. non-social versus social, and different types of social decisions).

Although our results did not statistically confirm a specific role of frontoparietal coherence for altruism in different inequality contexts (i.e., a significant interaction between entrainment type and inequality context), we did observe numerically stronger effects of gamma-band enhancement on altruistic choices (model-free) and preferences (model-based) during disadvantageous inequality. Viewed formally, these findings show that information sharing between other-interest signals in frontal regions and parietal evidence accumulation is necessary for altruism in general, irrespective of inequality context. However, the specific pattern that is visually evident in our data is clearly consistent with our findings that disadvantageous inequality is accompanied by stronger other-payoff EEG signals, and that fronto-parietal coherence correlates with altruistic preference stronger in this inequality context (Hu et al., 2023). This coherence between the previous EEG and the present tACS results suggests that frontoparietal coherence in the gamma-band may play a particularly marked role during disadvantageous inequality. While this proposition should be formally confirmed in future studies, it already now raises interesting questions about why putting individuals in a disadvantageous position should trigger particular motives that may rely on fronto-parietal coherence. One explanation may be attentional in nature: A disadvantageous position may make people more sensitive to the profit change of others who are already better off than themselves (Boyce et al., 2010; Payne et al., 2017), and this attentional sensitivity increase may correspond to a higher functional relevance of the oscillatory communication of the corresponding signals into the parietal evidence accumulation process. By contrast, participants in an advantageous position may have a more uniform decision weight on others’ profit and not be overly concerned about the specific profit change of the other person, as demonstrated by both our modeling results and by previous studies (Morishima et al., 2012). This could reduce individuals’ sensitivity to tACS entrainment of the corresponding signals into the evidence accumulation process, akin to a floor effect. Irrespective of such speculations, our results clearly inspire future studies to test more explicitly for differences in the role of frontoparietal coherence for different aspects of social decision-making that are thought to reflect different motives.

Taken together, our results demonstrate that increasing frontoparietal coherence by means of tACS can causally enhance individuals’ altruistic preferences and choices. This shows that altruistic choices have a clear neurobiological basis, which involves not just local neural computation of signals related to other’s outcomes but also functional integration of these signals with other (domain-general) areas involved in response preparation. Our findings thus also have novel implications for clinical neuroscience, as deficits in appropriate processing and integration of information about others usually lie at the core symptoms of several psychiatric and neurological disorders, including autism, psychopathy, and alexithymia (Alaerts et al., 2014; Contreras-rodríguez et al., 2014; FeldmanHall et al., 2013). Our approach may offer a promising start for developing intervention tools to improve individuals’ social functions in these populations.

## STAR Methods

### Participants

Forty-nine participants (20 females) participated in the study. Participants were informed about all aspects of the experiment and gave written informed consent. All the participants were right-handed, had normal or corrected-to-normal vision, were free of neurological or psychological disorders, and did not take medication during the time the experiment was conducted. They received between 109 – 133 CHF (depending on their choices) as compensation. Five participants were excluded from analyses because they failed to meet the requirements of the study (e.g., responding with both hands, not being able to stay awake during testing, no anatomical brain image available, and major in Psychology or Economy; remaining 44 participants: 19 – 32 years of age, mean 24.5 years). To determine the sample size, we followed our previous study and aimed to detect a medium effect with *d* = 0.4 (Hu et al., 2023). Analyses with G*Power 3.1 suggested that we need 47 participants to have a power of 1 – β > 0.85 to detect a medium effect with *d* = 0.4 at the level of α = 0.05 for one-tailed t-test. The behavioral results in the current study (i.e., t-tests of the gamma (vs sham or vs alpha) entrainment effect on the altruistic choice in the disadvantageous inequality context) showed a significant gamma entrainment effect with effect size *d* ≈ 0.38. Power analyses suggested that we can achieve a power of 1 – β ≈ 0.8 to detect this effect (*d* ≈ 0.38) with the remaining 44 participants at the level of α = 0.05 for one-tailed t-test. The experiment conformed to the Declaration of Helsinki, and the protocol was approved by the Ethics Committee of the Canton of Zurich.

### Behavioral paradigm

Each participant made 540 real decisions in a modified Dictator Game, requiring them to choose one of two allocations of monetary amounts between themselves and a receiver (another participant in the same study). We included two inequality contexts: In the disadvantageous context (DIS), the token amounts for participants were always lower than for the receiver; in the advantageous context (ADV), they were always higher. On each trial, one of the two options was revealed at the beginning of each trial (i.e., the reference option) and the other option was revealed at a later time (i.e., the alternative option, see below and Figure 1A for details). At the beginning of each trial, participants saw a central fixation cross together with a reference allocation option on one side of the cross. The alternative option, to be shown on the opposite side of the fixation cross, was initially hidden and replaced with “XX” symbols. After 0.5 s, the font color for all the stimuli changed from blue to green (the change direction of font color was counterbalanced across participants), indicating initiation of the trial. After a temporal jitter of 1 – 1.5 s (uniform distribution with a mean of 1.25 s), the alternative option was revealed (with XX symbols replaced with actual amounts). Participants had to choose the left or right option by pressing the corresponding keys on the response box with right index or middle finger within 2.5 s. The selected option was highlighted with color change. After another jitter of 0.5 – 2 s, the next trial started. Since all payoff stimuli were presented close to the fixation cross (i.e., visual angle smaller than 3°), participants were able to see the numbers clearly without shifting their gaze.

This sequential presentation allowed us to establish the inequality context with the presentation of the first option without having to explicitly instruct participants about the different contexts. There were 4 levels of reference options (i.e., 24/74, 24/98, 46/74, and 46/98) in each context. For the DIS context, the numbers before the slash denote the token amounts allocated to the participant and the numbers after the slash the token amount given to the partner, and vice versa for the ADV context. There were 22 or 23 alternative option levels corresponding to each reference option level; half of them were more equal than the reference option and the other half were more unequal than the reference option. The token differences for each party between the alternative and the reference options ranged from −19 to 19 (Fig. S1).

To avoid repetitions of exactly the same choices, we included three different trial sets with the same reference options and similar distributions of alternative options. The second and the third trial set were generated by adding a random jitter (i.e., −1, + 0, or +1) to self-/other-payoffs of the alternative options of the first trial set (Figure S1). In our trial set, we not only included decisions requiring opposite changes in self-and other-payoff (i.e., maximizing one and minimizing the other) but also a number of trials in which options lead to changes in inequality but increases or decreases for both self- and other-payoffs. However, the latter type of trial (number of trials = 208) was excluded from data analyses of altruistic choices, since participants’ choices in these trials cannot be unambiguously categorized as altruistic or selfish choices. Therefore, 332 out of 540 trials for each participant were kept in data analyses.

The task was divided into 6 runs, with each run lasting around 7 minutes. In each run, participants completed one block of the task (i.e., 30 trials, ∼130 s) with gamma entrainment, one block with alpha entrainment (i.e., 30 trials, ∼130 s), and two short blocks with sham stimulation (i.e., 15 trials, ∼83 s for each block). The order of the blocks in each run was counterbalanced across different runs, and the two sham stimulation blocks were always interleaved with gamma and alpha entrainment blocks (to prevent possible long-lasting effects due to two consecutive entrainment blocks). After each entrainment/stimulation block, participants rated their discomfort due to the stimulation in the current block on a Likert scale ranging from 0 to 20 (0 = not uncomfortable at all, 20 = very uncomfortable). After each run, participants had an approx. 1-min rest. In total, the task took between 50 to 60 minutes, including breaks between runs.

To avoid potential attentional or visual processing bias due to the fixed position of reference/alternative payoffs, we counterbalanced stimulus positions (left versus right) trial-by-trial within participant. To lower the processing load and avoid potential response errors due to misrecognition, we fixed the position of self/other payoffs within participants and counterbalanced it between participants.

Participation payment was determined at the end of the experiment and consisted of three parts: a base payment of 60 Swiss Francs (CHF, around $60 at the time of the experiment), one bonus (Bonus A) payment that depended on participants’ own decisions, and another bonus (Bonus B) payment that was determined based on the choices of previous participants whose outcomes were allocated to the current participant. To determine these bonus payments, the participant drew two envelopes from two piles of envelopes, one pile for bonus A and one pile for bonus B. Each envelope contained five different randomly determined trial numbers.

For Bonus A, the numbers in the envelope were the trial numbers that will be selected from the full list of the participant’s choices after the experiment. We calculated the mean payoff from these options and paid this as the first bonus.

For Bonus B, the numbers in the envelope were chosen from the full list of choices taken by previous participants. This list of choices was randomly drawn from the full list of choices of all previous participants, so that the participant was randomly paired with a different person on every round. The mean of the partner’s payoffs across the chosen five rounds was paid out as the second bonus. The final payments derived from the two bonuses were CHF 63.9 ± 5.3 (Mean ±SD).

The partner’s payoffs resulting from the participant’s own choices also entered the full list of choices for future participants, meaning that they were paid out to these participants if any of the current participant’s choices were selected. The exchange rate was 1 token = 0.5 CHF.

### Transcranial alternating current stimulation (tACS)

We delivered tACS entrainment by means of a multi-channel stimulator (DC-stimulator MC, NeuroConn Ilmenau, Germany). We first applied a topical anaesthetic crème (EMLA crème 5%) to the scalp location of the electrodes (Antal et al., 2017; Asamoah et al., 2019; van der Plas & Hanslmayr, 2020), which we removed after approximately 45 minutes to fix the tACS electrodes. This procedure helped improved blinding of the participants to the entrainment type (i.e. gamma tACS versus sham-tCS) and reduced the skin sensations due to tACS, thereby making the entrainment more comfortable (McFadden et al., 2011).

Following our previous EEG study, we placed two arrays of small electrodes (3 × 1 electrode montage: 3 peripheral electrodes and 1 central electrode, electrode diameter: 2 cm, electrode center-to-center distance: 5 cm) to focally stimulate the frontal and parietal regions over which we had observed interregional phase coupling to correlate with altruistic behavior during disadvantageous inequality. The central electrodes were placed at the centers of the significant clusters (channel no. 37, 39, 61, 65, and 70 for the parietal region and channel no. 15, 19, 24, 28, 69, 71, 86, 87, 106, 107, 108, 112, and 113 for the frontal region on 128 channel waveguard cap, ANT Neuro, Netherlands) identified in our previous EEG study (Hu et al., 2023). We used a conductive paste (Ten20 EEG Conductive Paste, Weaver and Company, Colorado, USA) to fixate the electrodes on the scalp and delivered alternating currents at 72 Hz (gamma band), modulated with a 6Hz envelope to approximate the phase-amplitude modulation occurring endogenously in the human brain (Canolty et al., 2006). We also included a control entrainment condition in which we delivered alternating currents at 12 Hz (alpha band) modulated with a 6 Hz envelope. For the sham stimulation condition, we delivered alternating currents as in the gamma or alpha entrainment conditions, but only for 2 s (1-s ramp-up and 1-s ramp-down phases) every 27 s during the block. The maximum peak-to-peak current intensity ranged from 2.4 mA to 4 mA, depending on each participant’s tolerance for the stimulation currents (tested before the experimental runs were conducted). Participants were not aware of which type of entrainment they received in each block, or what hypotheses about the effects of entrainment on their behavior were tested.

We estimated the electric field for our tACS electrode montage by means of the Simnibs 2.1 toolbox (Makarov et al., 2019). The estimated electric field clearly showed that the targeted parietal and frontal areas under the two main tACS electrodes are affected by the stimulation with a relatively good special focality (see Figure 1B).

### Behavioral analysis

To examine the effect of tACS on altruistic behaviors, we performed logistic mixed-effects regression of individuals’ choices (altruistic = 1, selfish = 0) on various regressors of interest, including inequality context (DIS = 0, ADV = 1), gamma entrainment (gamma = 1, sham or alpha = 0), sham stimulation (sham = 1, gamma or alpha = 0) or alpha entrainment (alpha = 1, sham or gamma = 0), and the interactions between inequality context and entrainment type. To test whether or not physical discomfort related to different tACS entrainment types and current intensity affected participants’ choices, we also conducted regression models with each participant’s discomfort rating for different types of entrainment and the individual current intensity as regressors of no interest. All mixed-effects regression models included random effects for the participant-specific intercept term. The results of these regressions are reported in Table 1. We performed these mixed-effects regression analyses using the lme4 package in R.

Since our previous study had shown that the ADV context enhances altruistic behavior relative to the disadvantageous context, and that greater gamma-band phase coupling is associated with stronger altruistic preferences, we tested the directional hypotheses corresponding to these effects in mixed-effects regression models with one-tailed statistical tests. Specifically, in the regression models reported in Table 1 and Table S1, S2, and S4, we applied one-tailed statistical tests for the effects of “Inequality context” and “Gamma” entrainment, but two-tailed statistics for the other effects with undirected hypotheses. Likewise, in the post-hoc analyses contrasting altruistic choices and parameters between different conditions, we applied one-tailed t-test statistics with Bonferroni correction for multiple comparisons.

### Computational model

In the first model (M1), the subjective value difference between the more generous option and the more selfish option (VD) is constructed based on the Charness-Rabin utility model (Charness & Rabin, 2002)(Charness & Rabin, 2002)(Charness & Rabin, 2002) in which people assign complementary weights to outcomes for themselves or others:

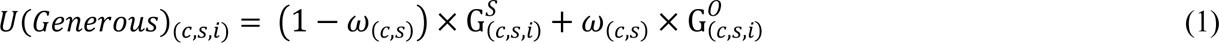

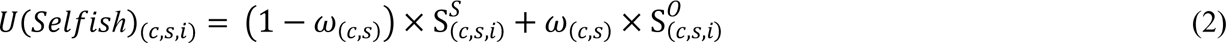

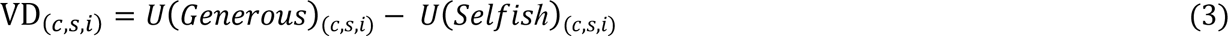

with indices c for conditions (c = DIS & Gamma, DIS & Sham, DIS & Alpha for different entrainment conditions in disadvantageous inequality context, and c = DIS & Gamma, DIS & Sham, DIS & Alpha for different entrainment conditions in advantageous inequality context), s for participants (s = 1, …, Nparticipants), and i for trials (i = 1, …, Ntrials). 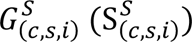 indicates participants’ payoff of the generous (selfish) option in condition c, for participant s and trial i; 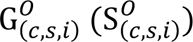 indicates the partners’ payoff in the generous (selfish) option in condition c, for participant s and trial i.

We calculated trial-by-trial value difference (VD_(*c*,*s*,*i*)_) between the two offers (VD_(*c*,*s*,*i*)_ = *U*(*Generouss*)_(*c*,*s*,*i*)_ − *U*(*Selfish*)_(*c*,*s*,*i*)_) and employed a softmax function to transform these value differences into probabilities of choosing the more generous option:

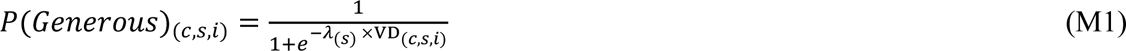

where *λ*_(*s*)_ is participant-specific inverse temperature parameters reflecting to what extent individuals’ decision depend on VD.

In the second model (M2), we included condition-specific inverse temperature parameters to examine the extent to which individuals’ decision weigh on value difference across different conditions:

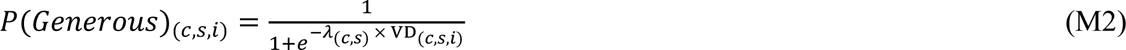

where *λ*_(c,*s*)_is condition-specific free temperature parameters reflecting individuals’ weight on value difference. The calculation of value difference (VD_(*c*,*s*,*i*)_) in M2 was the same as M1.

In the third model (M3), we included a third set of parameters (*μ*_(c,*s*)_) to measure to what extent individuals’ decisions weigh on the overall benefit for all parties (i.e., efficiency):

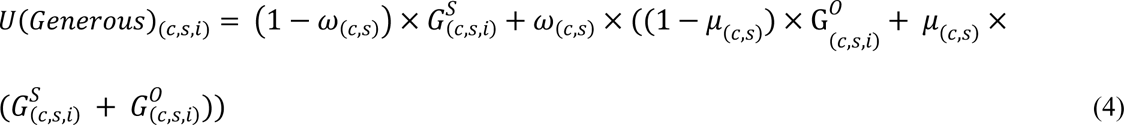

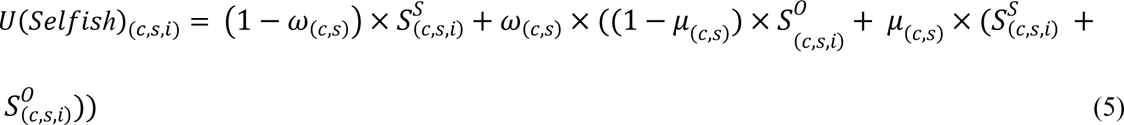

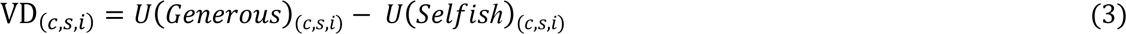

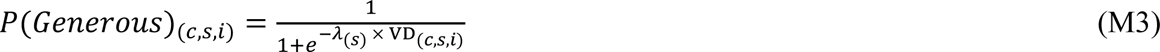

where *μ*_(c,*s*)_ is the condition-specific parameters measuring individuals’ weight on efficiency and *λ*_(*s*)_ is the participant-specific inverse temperature parameters.

We obtained best fitting parameters by maximizing the log-likelihood of the data for each model with the MATLAB function fmincon. To avoid the optimization getting stuck in local minima, we used 50 multiple starting points. To evaluate model fits, we calculated the Bayesian Information Criterion (BIC) (Schwarz, 1978) which rewards model parsimony to avoid overfitting:

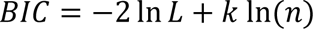

where *L* is the maximized likelihood for the model, *k* is the number of free parameters in the model, and *n* is the number of observations.

We also followed established procedures (Lewandowsky & Farrell, 2011) to calculate Bayes factor as BF = exp(− 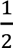 ΔBIC), where Δ*BIC* is the difference in the average BIC between the winning model (M1) and each alternative model. BF between 3 and 10 indicates moderate evidence, BF > 10 indicates strong evidence, and BF > 100 indicates very strong evidence that the winning model is superior to the alternative model (Lewandowsky & Farrell, 2011).

### Parameter recovery

We used the fitted parameters from each individual participant to simulate choices with the winning model (M1) 100 times, fitted the model to these simulated choices by maximizing the log-likelihood of the data with the MATLAB function fmincon, and correlated the averaged recovered parameters (of the 100 simulations) with the ones used to simulate choices. All parameters recovered well (Figure S3, correlations between simulated and recovered parameters p < 0.001 for all parameters).

---

**Table S1.** Logistic mixed-effects model results of choice data for DIS context.

| Fixed effects | Model 1 |  | Model 2 |  | Model 3 |  | Model 4 |  |
| --- | --- | --- | --- | --- | --- | --- | --- | --- |
| | $\beta$<br>(95% CI) | p-value | $\beta$<br>(95% CI) | p-value | $\beta$<br>(95% CI) | p-value | $\beta$<br>(95% CI) | p-value |
| Intercept | -2.70***<br>(-3.11 – -2.30) | < 0.001 | -2.71***<br>(-3.12 – -2.30) | < 0.001 | -2.73***<br>(-3.14 – -2.32) | < 0.001 | -2.69***<br>(-3.10 – -2.28) | < 0.001 |
| Gamma (G) | 0.18*<br>(-0.01 – 0.37) | 0.035 | 0.19*<br>(-0.01 – 0.38) | 0.029 | 0.21 *<br>(-0.001 – 0.42) | 0.025 | 0.17*<br>(-0.03 – 0.37) | 0.046 |
| Sham (S) | -0.01<br>(-0.21 – 0.19) | 0.929 | - | - | 0.04<br>(-0.20 – 0.27) | 0.754 | - | - |
| Alpha (A) | - | - | 0.01<br>(-0.19 – 0.21) | 0.929 | - | - | -0.04<br>(-0.27 – 0.20) | 0.754 |
| Discomfort rating | - | - | - | - | 0.08<br>(-0.13 – 0.29) | 0.469 | 0.08<br>(-0.13 – 0.29) | 0.469 |
| Intensity | - | - | - | - | 0.01<br>(-0.37 – 0.40) | 0.951 | 0.01<br>(-0.37 – 0.40) | 0.951 |
| Conditional R<sup>2</sup> | 0.32 |  | 0.32 |  | 0.32 |  | 0.32 |  |
| LL | -2078 |  | -2078 |  | -2077 |  | -2077 |  |
| BIC | 4191 |  | 4191 |  | 4208 |  | 4208 |  |

Tests for the effects were implemented with one-tailed statistical tests. Discomfort rating, participants rated their discomfort due to the stimulation after each entrainment/stimulation block. Intensity, each participant was stimulated with an electric current intensity on his/her tolerance level for the stimulation currents tested before the experiment runs. Gamma (G): gamma entrainment; Sham (S): sham stimulation; Alpha (A): alpha entrainment; LL: log-likelihood; BIC: Bayesian Information Criterion. \*\*\*, $p < 0.001$ ; \*\*, $p < 0.01$ ; \*, $p < 0.05$ .

**Table S2.**
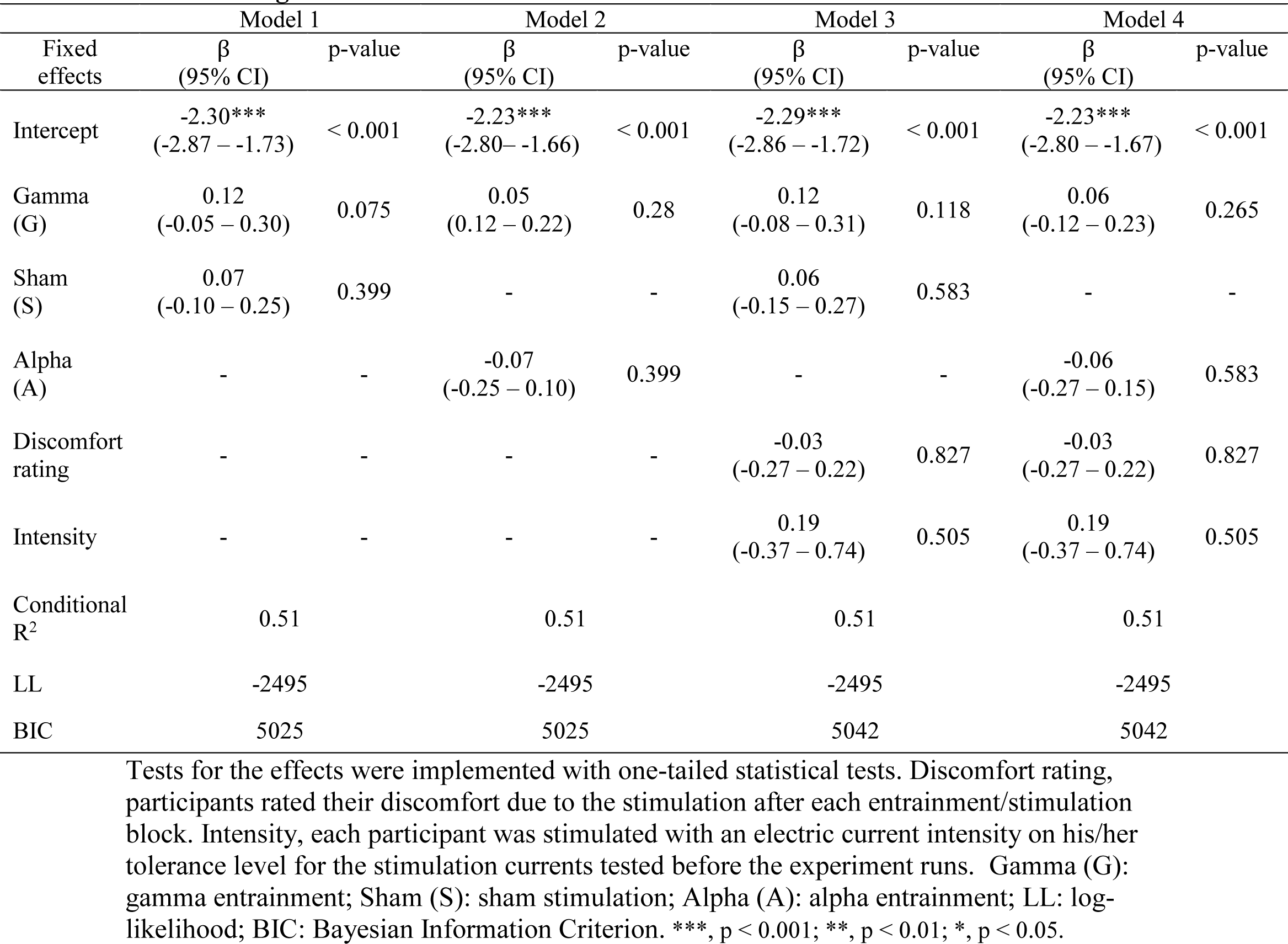
Logistic mixed-effects model results of choice data for ADV context.

**Table S3.** Model comparison results.

| Model | Parameters | BIC (mean ± SE) | BF |
| --- | --- | --- | --- |
| M1 | $\omega_{(c,s)}, \lambda_{(s)}$ | $202 \pm 14$ | 1 |
| M2 | $\omega_{(c,s)}, \lambda_{(c,s)}$ | $218 \pm 15$ | $2.98 \times 10^3$ |
| M3 | $\omega_{(c,s)}, \mu_{(c,s)}, \lambda_{(s)}$ | $236 \pm 14$ | $2.42 \times 10^7$ |

BIC, Bayesian Information Criterion; BF, Bayes factor. $\omega_{(c,s)}$ , relative weight on others' payoffs; $\lambda_{(c,s)}$ , inverse temperature parameter; $\mu_{(c,s)}$ , relative weight on efficiency; c for conditions (c = DIS & Gamma, DIS & Sham, DIS & Alpha for different stimulation conditions in disadvantageous inequality context, and c = DIS & Gamma, DIS & Sham, DIS & Alpha for different stimulation conditions in advantageous inequality context), s for participants (s = 1, ..., N<sub>participants</sub>). SE, standard error.

**Table S4.** Logistic mixed-effects model results of simulated choice data.

| Fixed effects | Model 1 |  | Model 2 |  |
| --- | --- | --- | --- | --- |
| | $\beta$<br>(95% CI) | p-value | $\beta$<br>(95% CI) | p-value |
| Intercept | -2.71***<br>(-3.13 – -2.29) | < 0.001 | -2.71***<br>(-3.13 – -2.29) | < 0.001 |
| Inequality context (C) | 0.84***<br>(0.66 – 1.02) | < 0.001 | 0.94***<br>(0.76 – 1.11) | < 0.001 |
| Gamma (G) | 0.23**<br>(0.04 – 0.42) | 0.008 | 0.24**<br>(0.05 – 0.43) | 0.007 |
| Sham (S) | -0.008<br>(-0.20 – 0.19) | 0.937 | - | - |
| Alpha (A) | - | - | 0.01<br>(-0.19 – 0.20) | 0.937 |
| G*C | -0.12<br>(-0.37 – 0.12) | 0.324 | -0.22*<br>(-0.47 – 0.03) | 0.081 |
| S*C | 0.10<br>(-0.16 – 0.35) | 0.459 | - | - |
| A*C | - | - | -0.10<br>(-0.35 – 0.16) | 0.459 |
| Conditional R<sup>2</sup> | 0.37 |  | 0.37 |  |
| LL | -4961 |  | -4961 |  |
| BIC | 9988 |  | 9988 |  |

Tests for the effects were implemented with one-tail statistics. Gamma (G): gamma entrainment; Sham (S): sham stimulation; Alpha (A): alpha entrainment; LL: log-likelihood; BIC: Bayesian Information Criterion. \*\*\*, $p < 0.001$ ; \*\*, $p < 0.01$ ; \*, $p < 0.05$ .

**Figure S1.**
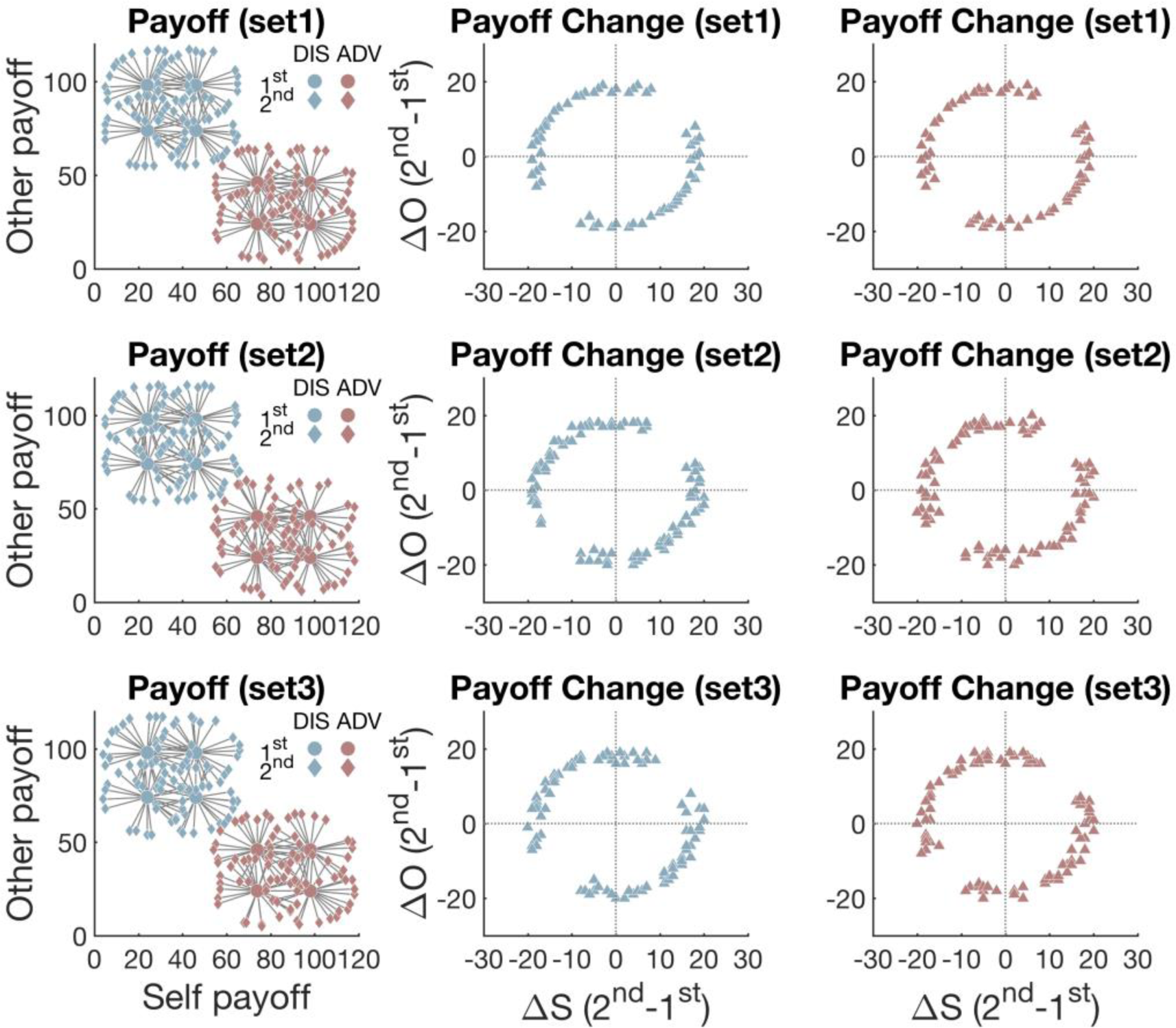
Payoff schedule. We included two inequality contexts: disadvantageous inequality (DIS) and advantageous inequality (ADV). In the left panel, each dot represents one allocation option and each gray line represents one pair of options that was presented to participants. Blue dots are options in DIS and pink dots are options in ADV. Dots in the center of the circle are the 1^st^ options and diamond dots are the 2^nd^ options. Middle and right panels show the distributions of the self-/other-payoff changes between the 2^nd^ and the 1^st^ option (ΔS and ΔO) in DIS and ADV, respectively. These three sets of payoff matrices (top, middle, and bottom panel) have the same reference options and similar distributions of alternative options. By having such payoff matrices of all trials, we matched self-/other-payoff differences and the resulting absolute levels of inequality across both contexts and also across the 2^nd^ and the 1^st^ options. This allowed us to compare choices and response times, as well as neural processing of different choice features (self- and other-payoff, inequality), between the two contexts.

**Figure S2.**
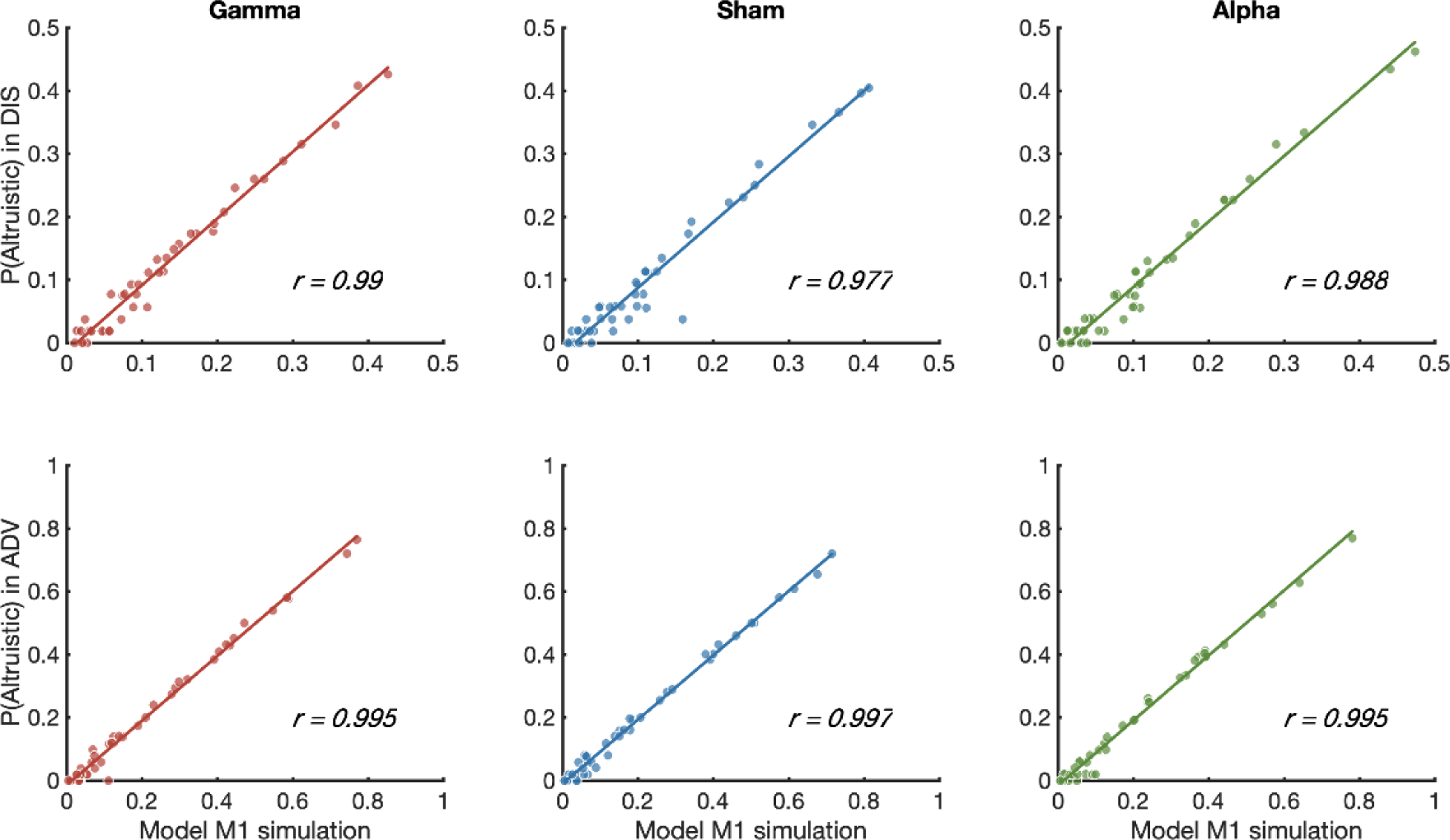
Model simulation results. Correlations of the probability of altruistic choice between participants’ true responses (y axis) and model M1 simulation responses (x axis) based on the winning model, for DIS context (A) and for ADV context (B). Model simulation data are highly correlated with observed true data across all inequality contexts and stimulation types.

**Figure S3.**
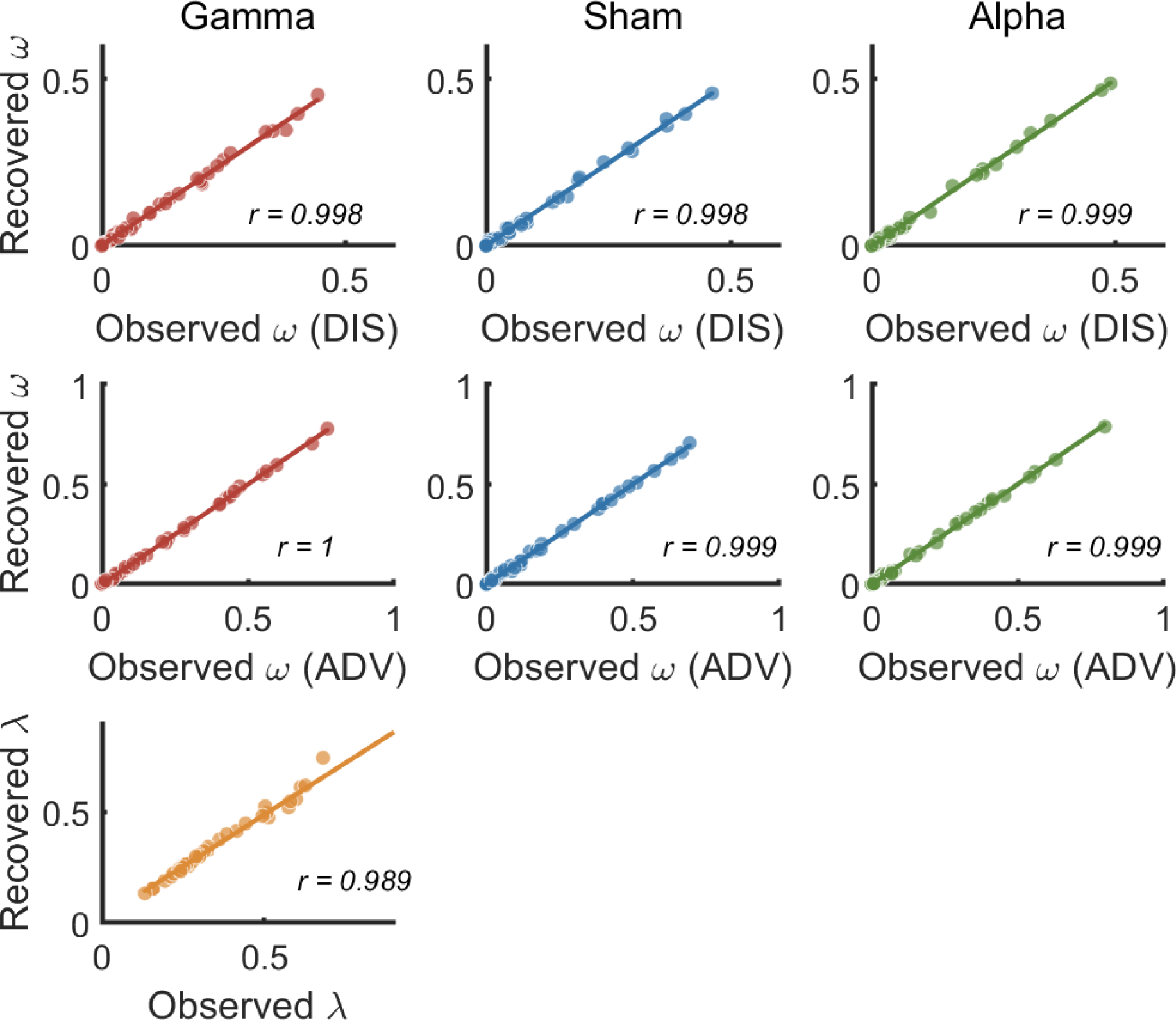
Parameter recovery. Observed parameters (values fitted to participants’ choice data in the winning model M1) are used to simulate choices, and these choices are used to recover the observed parameters. We repeated this procedure 100 times to obtain the average value of each recovered parameter and correlated the averaged recovered parameters with the observed parameters (which were used to generate the choices). Each dot plots the averaged recovered parameter from the simulated choices against the observed parameter (generating parameter), and the colored lines represent the regression fits of the observed parameters on the recovered parameters.

